# Transcription Factor Driven Gene Regulation in Autism Spectrum Disorder

**DOI:** 10.1101/2025.02.24.639835

**Authors:** Nimisha Ghosh, Walter Arancio, Tariq Al Jabry, Raya Al-Maskari, Daniele Santoni

**Affiliations:** Department of Computer Science and Engineering, Shiv Nadar University, Chennai, Tamil Nadu, India; Institute for System Analysis and Computer Science “Antonio Ruberti”, National Research Council of Italy, Via dei Taurini 19, Rome, 00185, Italy; Institute for Research and Biomedical Innovation, National Research Council of Italy, Via Ugo la Malfa 153, Palermo, 90146, Italy; College of Medicine and Health Sciences, Sultan Qaboos University, P.O. Box 35, Al Khod, 123, Oman

**Keywords:** Autism Spectrum Disorder, Transcription Factors, gene expression regulation, Pearson Correlation

## Abstract

Autism Spectrum Disorders (ASD) encompass a group of neurodevelopmental disorders in which an affected individual faces challenges in social interaction and communication, along with restricted and repetitive stereotypic behavioral patterns and interests. In this work, we have studied the differential gene regulation between patients and controls, mediated by Transcription Factors (TFs), of key genes involved in ASD. Nine and seven TFs have been identified as potential regulators of the set of syndromic and non-syndromic key high confident genes retrieved by the Simons Foundation Autism Research Initiative (SFARI) database. We have also identified significant couples of Transcription Factor - Target Gene potentially associated with an altered regulation in ASD patients. Consistently, many identified couples are involved in processes associated with brain morphogenesis and development. In this regard, this biased regulation could be the target of some experimental design in order to 1) test this hypothesis and 2) try to target this altered regulation pattern in ASD samples. In conclusion, we would like to emphasize that the present work proposes an effective and reliable computational approach that could be applied to any disease with known key genes and available gene expression data.

## 1 Introduction

Autism Spectrum Disorders (ASDs) encompass a group of neurodevelopmental disorders in which an affected individual faces challenges in social interaction and communication, along with restricted and repetitive stereotypic behavioral patterns and interests [1]. Individuals with autism can be diagnosed as early as 2-3 years, while the mean age of diagnosis is 4-5 years [2]. It should be noted that more adults are being evaluated for possible cases of autism. According to data from CDC’s Autism and Developmental Disabilities Monitoring (ADDM) Network, an average of 1 in every 36 (2.8%) 8-year-old child was estimated to have ASD in 2020. ASD is reported to occur in all ethnic groups, with a prevalence of 3.8 times higher among boys (4.3%) compared to girls (1.1%) [3]. The etiology of ASD is likely to be multifactorial, with both genetic and non-genetic factors playing a role in its onset. This complexity is well represented by the differentiation within ASD between syndromic and non-syndromic forms [4]. While the syndromic ASD etiology is well defined and it seems to be often associated with chromosomal abnormalities or monogenic alterations, the non-syndromic ASD is still relatively undefined. A collaboration between de novo mutation and prenatal and postnatal environmental factors appears to be playing a major role in its onset. However, ASD is considered a highly heritable genetic disorder, with genome-wide association studies (GWAS) and next-generation sequencing identifying numerous risk genes. In 20-25% of ASD cases, the genetic basis can be attributed to de novo mutations, rare and common genetic variants, or ASD-associated polymorphisms [5]. The functional implications of these genetic associations remain poorly understood, necessitating integrative analyses that connect genetic risk with regulatory mechanisms.

The Simons Foundation Autism Research Initiative (SFARI) database, a curated resource for ASD research, catalogs genes associated with ASD and scores them based on their relevance to autism risk ^1^. This database provides a foundation for studying gene expression regulation, which includes Transcription Factors (TFs) as well. Transcription Factors (TFs) are a group of proteins that can regulate gene expression [6]. The study of TFs is thus important to understand the regulation of genes as well as their differential role in ASD patients and healthy controls.

In this regard, Harris et al. [7] have considered 38 individuals with variants in RFX family of genes (RFX3, RFX4 and RFX7) where RFX are evolutionary conserved transcription factors acting as master regulators of ciliogenesis and central nervous system development. The authors presented the analysis of the expression patterns and the downstream analysis of the targets of the aforementioned genes in order to relate them to other neurodevelopmental risk genes. This study revealed that the 38 individuals share traits of ASD. Another study by Pastore and colleagues [8] has shown that a specific risk factor for ASD is PTCHD1 which has both variable temporal and brain subregion-specific levels of transcription in embryonic and postnatal development of mice. Moreover, the single nucleotide polymorphism (SNP) rs7052177 is also found to be related to ASD [9]. This SNP has further association with TFs such as STAT3, STAT5A and STAT5B which affect the expression of PTCHD1 [9]. In [10], Ahmad et al. have looked into the role of CD45 signaling in children with ASD. In this regard, they have investigated the role of CD45 cells expressing inflammatory transcription factors in ASD. Their results show that CD45 has the possibility of playing an important role in the immune abnormalities of ASD. Binding sites of specificity protein 1 (SP1) have been found in the promoter regions of many genes involved in autism. Considering this fact, [11] has hypothesised that the dysfunction of SP1 can affect the expression of candidate genes of multiple autism, thereby contributing to the heterogeneity of autism. In their work, they have observed the higher expression of SP1 in (anterior cingulate gyrus (ACG)) of autism patients as opposed to healthy controls. The authors in [12] have considered transcription factor SP9 to study ASD and other intellectual disability. Their study provides both *in silico* and *in vitro* evidence of the presence of novel form of interneuronopathy caused by de novo heterozygous variants in SP9. In [13], the authors have focussed on FOXP family of TFs due to the evidence that such genes are linked to ASD or other genes influencing ASD. There are three genes in the FOXP family that are expressed in central nervous system, FOXP1, FOXP2 and FOXP4. Such TFs play major role in brain development and language evolution as well. Coskunpinar et al. [14] have focussed on the variant changes in Gamma amino butyric acid (GABA) receptor subunit genes (GABRB3, GABRG3 and the HTR2A) encoding one of the serotonin receptors. They have conducted their study considering 200 patients with ASD between the ages of 3-9 and 100 healthy controls where the DNA isolation has been performed from peripheral blood samples. Their study has shown that for HTR2A and GABRG3, the homozygous C genotype and homogyzous T genotype respectively are much higher in patients than in controls, thereby confirming their role in the predisposition of ASD. To understand the role of TCF7L2 in the pathogenesis in ASD, Hryniewiecka et al. [15] have conducted experiments to investigate the functional consequences of the deficiency of TCF7L2 in the brain. From their study, they have drawn the conclusion that the disruption of thalamic locus of TCF7L2 may be crucial for autistic symptoms development.

In the present work we propose a method to reveal those TFs responsible for a differential regulation of some of their target genes in ASD with respect to controls. In Materials and Methods section we first explain how we have selected the genes involved in ASD, both syndromic and non-syndromic, through SFARI database. Then we describe the software TRANSPARENT [16] that we have used to identify TFs potentially able to regulate the set of considered genes. We finally introduce the gene expression data of ASD and Controls we have used in this work. In the Result section, we present and discuss the identified TFs, the Transcription Factors - Target Genes couples and analyse the Pearson correlation values using gene expression data from ASD and controls. Finally, we report those couples that show a differential correlation between ASD and Controls, hence a differential gene regulation between patients and controls.

## 2 Materials and Methods

The experiments in this work are carried out according to the pipeline as given in Fig. 1.

**Fig. 1.**
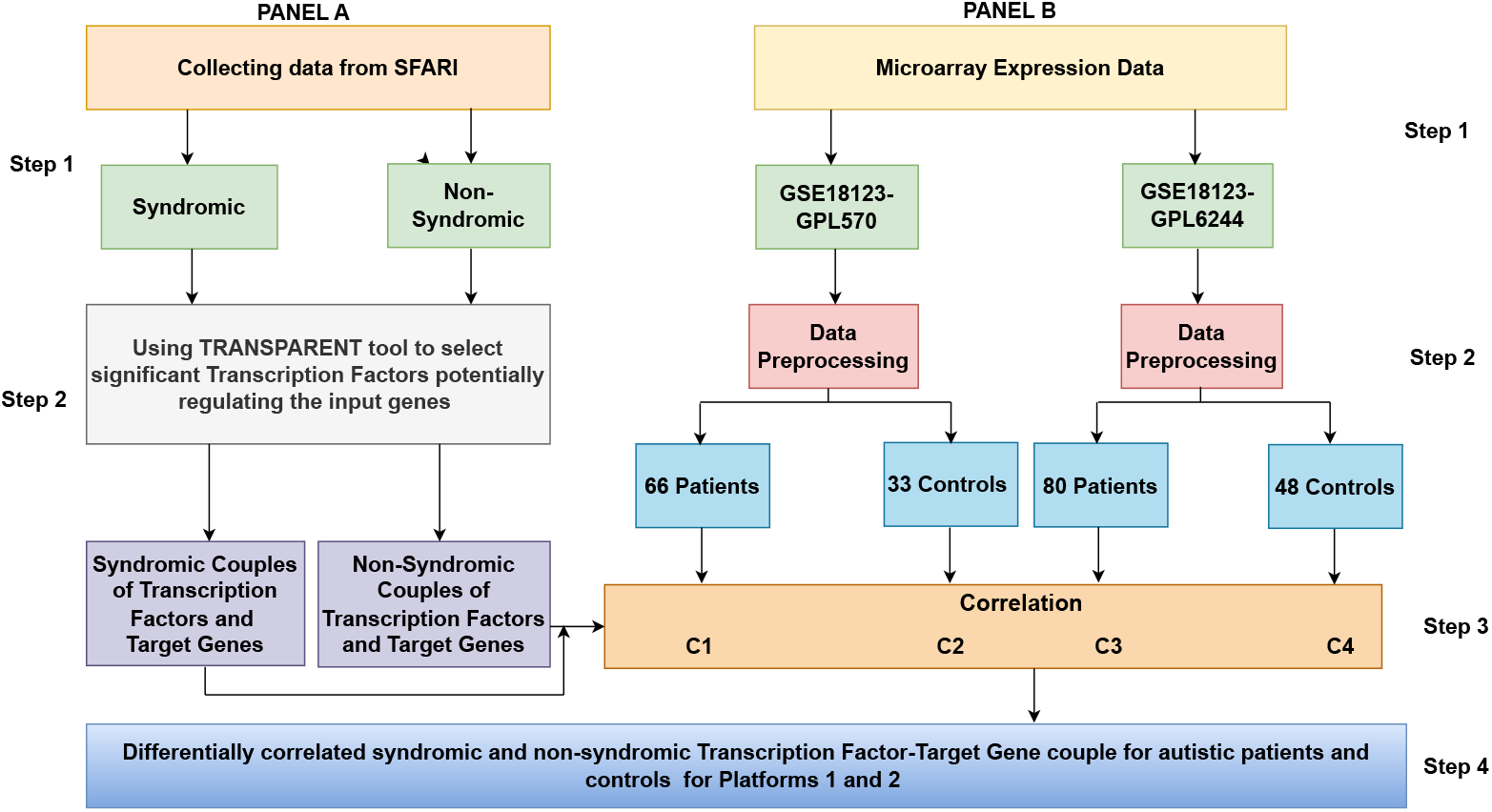
Pipeline of the work

Initially, as part of data collection in Step 1A, genes associated with ASD (ASD) were retrieved from SFARI database ^2^. Those genes with high confidence score (level S1 in the database) indicating a strong reliable association with ASD were selected. They were further divided into two categories with 116 syndromic and 117 non-syndromic genes. The list of genes for both the categories is available as Supplementary Material STAB1. In Step 2A, TRANSPARENT tool [16] was used to select significant TFs potentially regulating the two lists of syndromic and non-syndromic genes. This led to the identification of TF - target gene couples for such two lists. In Step 1B of the analysis, gene expression microarrays of autistic patients and controls were collected from GEO database (Accession 18123) platforms GPL570 and GPL6244. The expression data were extracted from blood of ASD patients and controls. In this regard, GPL570 has 66 male patients and 33 male controls while GPL6244 has 104 patients (80 male and 24 female) and 82 controls (48 male and 34 female). Since significant sex- and gender-specific characteristics and differences in the biology of autism have been observed [17], we decided to conduct the analysis exclusively on male samples, excluding female ones. In Step 2B data preprocessing was carried out where a single expression value for each given gene was obtained by averaging values of all the probes associated with this gene. Furthermore, computed expression values were normalised in log exponential scale. Subsequently, such data for autistic patients and controls were separately analysed. In Step 3, Pearson correlation analysis between expression values of the TF-target gene couples was carried out as identified in Step 2A of the pipeline. The correlation was computed for both GPL570 and GPL6244 considering separately ASD patients and controls. This analysis was carried out for both syndromic and non-syndromic TF-target gene couples. P-values were computed through Pearson correlation and then adjusted based on Benjamin-Hochberg correction for multiple hypothesis testing. This led to the calculation of 4 sets of P-values for syndromic and 4 for non-syndromic: Syndromic couples for patients and controls for GPL570 and GPL6244 and similarly for non-syndromic couples as well.

## 3 Results

In this Section we present results obtained by following the analysis as described in Fig.1. We first show the list of TFs (see Table 1), potentially involved in the regulation of syndromic and non-syndromic genes, obtained through TRANSPARENT software. We then analyse Pearson correlations between identified TFs and their target genes and we finally select, according to the adjusted P-values, those couples showing a differential correlation between ASD and Controls (see Table 2).

**Table 1.** Transcription Factors identified through TRANSPARENT potentially regulating the genes of the considered set (Syndromic/Non Syndromic). The Gene Symbol is reported in the first column while the gene set either S (Syndromic) or NS (Non Syndromic) is reported in the second column. The adjusted P-value, as provided by TRANSPARENT, the number of target genes TG (those genes showing at least one predicted TFBS in their promoter region) and a brief description of the TF function are reported in columns three, four and five respectively.

| TF | S/NS | Adj P-value | TG | Description |
| --- | --- | --- | --- | --- |
| KLF15 | Syn | 2.52E-11 | 94 | Kruppel-like factor 15 |
| **KLF4** | Syn | 9.69E-09 | 96 | Kruppel-like factor 4 |
| **VEZF1** | Syn | 3.65E-07 | 93 | Vascular Endothelial Zinc Finger 1 |
| **ZNF148** | Syn | 4.62E-06 | 79 | Zinc Finger Protein 148 |
| SP9 | Syn | 3.19E-05 | 94 | Specificity Protein 9 |
| FOXP2 | Syn | 0.00030 | 63 | Forkhead Box P2 |
| **MZF1** | Syn | 0.00035 | 88 | Myeloid Zinc Finger 1 |
| YY2 | Syn | 0.00044 | 37 | Yin Yang 2 |
| MAZ | Syn | 0.00048 | 90 | Myc-associated zinc finger protein |
| **VEZF1** | Non-Syn | 2.95e-06 | 95 | Vascular Endothelial Zinc Finger 1 |
| ZBTB14 | Non-Syn | 4.65E-06 | 38 | Zinc Finger and BTB Domain Containing 14 |
| ZNF263 | Non-Syn | 1.65E-05 | 90 | Zinc Finger Protein 263 |
| **MZF1** | Non-Syn | 8.55E-05 | 90 | Myeloid Zinc Finger 1 |
| **ZNF148** | Non-Syn | 0.00018 | 76 | Zinc Finger Protein 148 |
| SOX14 | Non-Syn | 0.00023 | 18 | SRY-box transcription factor 14 |
| **KLF4** | Non-Syn | 0.00037 | 88 | Kruppel-like factor 4 |

**Table 2.**
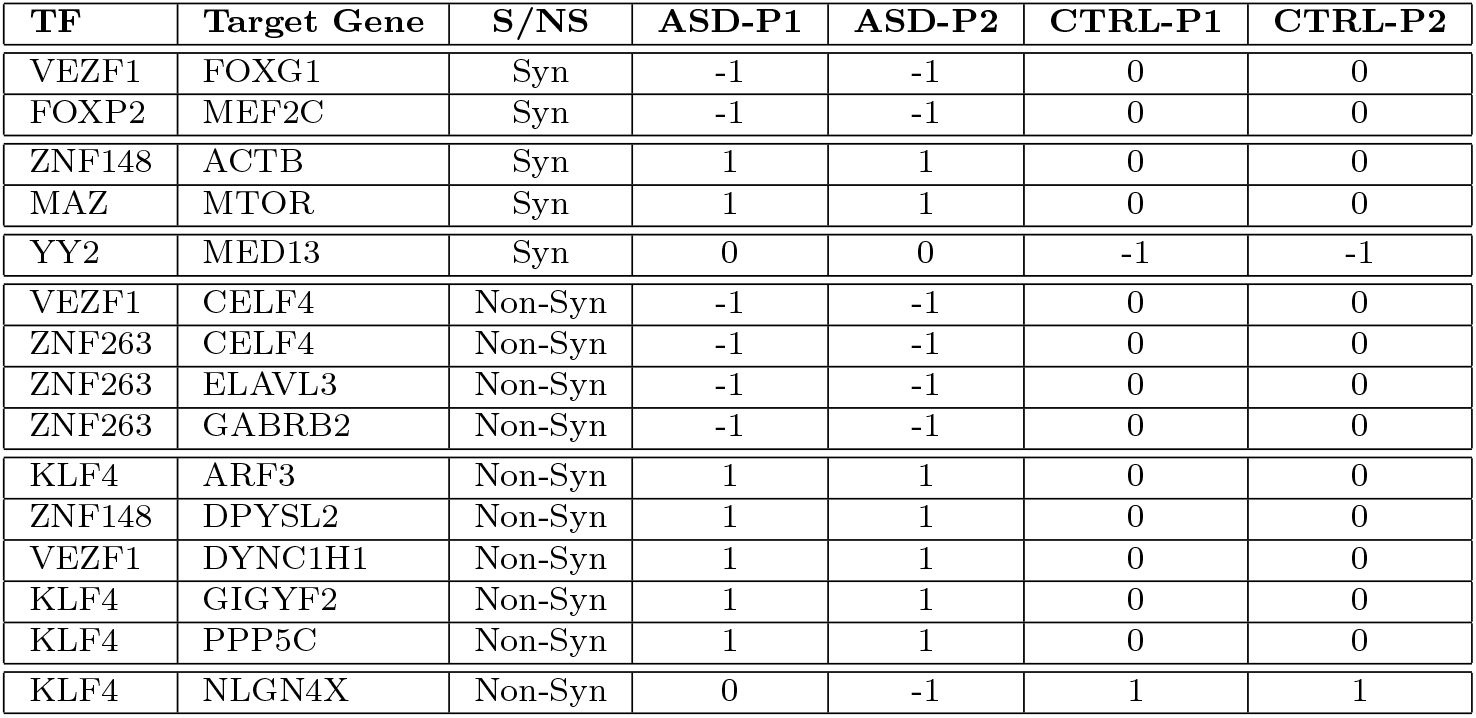
Differential Pearson Correlation between expression values of Transcription Factor (column1) and Target Gene (column2). P1 and P2 refer to as GPL570 and GPL6244 respectively. CTRL refers to as Healthy Controls and ASD to Patients affected by Autistic Spectrum Disease.

### 3.1 Selecting Transcription Factors regulating Syndromic and Non-Syndromic ASD genes

Starting from 116 syndromic and 117 non-syndromic genes which was obtained from SFARI database as explained in 2, we identified those TFs that are likely to be involved in the regulation of these genes. We used TRANSPARENT tool [16] to extract those significant TFs separately for syndromic and non-syndromic. Each identified TF shows a significant Transcription Factor Binding Site (TFBS) enrichement (P-value *<* 0.01) in the promoter regions of the considered genes. The list of the selected TFs is reported in Table 1 along with the set of genes (syndromic or non-syndromic) they are supposed to regulate, the adjusted P-value, the number of target genes (those genes showing at least one predicted TFBS in their promoter region) and brief description of the TF function. Setting a threshold for P-value *<* 0.01 7 TFs were identified for non-syndromic genes and 9 for syndromic ones. Significantly even if the analyses were performed on two different promoter sequence sets (with no gene in common), 4 TFs (highlighted in the table in bold) occur in both lists VEZF1, MZF1, ZNF148 and KLF4 providing consistency and reliability to the analysis and suggesting a relevant likelihood of the involvement of those TFs in ASD. The obtained set of TFs is also consistent with results shown in [16] where the analysis was performed starting from a much larger set of 1112 genes associated to autism.

### 3.2 Analysis of significantly correlated Transcription Factor - Target Gene couples using expression data

We first identified all the couples made up of a significant TF, as reported in Table 1, and one of its target genes (a gene whose promoter region contains at least one TFBS of the given TF, the number of such genes is reported in the TF column in Table 1). In this regard, two lists of TF-target gene couples were obtained; the former for syndromic and the latter for non-syndromic. For each given couple (both for syndromic and non-syndromic) we computed the Pearson correlation value separately for ASD and control samples using data values coming from both GPL570 and GPL6244. For each couple, we obtained four correlation values (see Fig.1 Panel B - Step 3): two for ASD (GPL570 and GPL6244 - C1 and C3 respectively) and two for controls (GPL570 and GPL6244 - C2 and C4 respectively). All P-values associated with the Pearson correlations were adjusted for multiple tests applying Benjamini Hochberg correction. We considered two P-value thresholds: one more strict equal to 0.01 and one less strict equal to 0.05. For each considered couple and each dataset, out of the four, we refer to *C* and *P* as the Pearson correlation and the adjusted P-value respectively. We associated a value *S*(*C, P*) equal to −1, 0, 1 or *N* with each C,P couple according to the following rule:

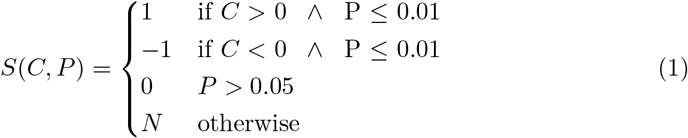

We assigned 1 or -1 when *P <*0.01 but we assigned 0 when *P >*0.05 instead of *P* 0.01 to set a stricter constraint.

### 3.3 Identification of differential gene regulation between ASD and controls mediated by Transcription Factors

Formula 1 was designed to easily and effectively identify those couples that show a significantly differential correlation between ASD and Healthy samples (Step 4). For example, if for a given couple TF (Transcription Factor) - G (target gene), S(C, P) is equal to +1 (or -1) in ASD (for both PL570 and GPL6244) and equal to 0 in controls (for both PL570 and GPL6244) we can conclude that the expression of G is differentially regulated by TF in ASD and Healthy samples.

We identified TF-G couples potentially involved in altered expression regulation in ASD for both syndromic and non-syndromic as reported in Table 2. If a statistically significant negative regulation is observed between a given TF and one of its target genes in patients but not in controls, this differential regulation can be a cause and/or consequence of the disease and thus it can be further studied. This is the case of VEZF1 that negatively regulates FOXG1 in ASD patients while no correlation occurs in Controls. In Fig. 2 VEZF1 expression values (x-axis) are plotted against FOXG1 expression values (y-axis) for ASD (red circles) and Controls (green triangles). Panel A reports expression values collected from GPL570 while panel B reports expression values collected from GPL6244. A negative significant Pearson correlation can be observed for ASD in both panels (P-value equal to 0.0032 and 0.0005 for GPL570 and GPL6244 respectively) where the linear regression (m=-0.72 and m=-1.43) is also reported (red lines). In both panels the green triangles, representing Controls, highlight no significant correlation between VEZF1 ansd FOXG1 (P-value equal to 0.1357 and 0.0935 for GPL570 and GPL 6244 respectively) suggesting that the negative regulation mechanism occurring in ASD samples is not active in Controls.

**Fig. 2.**
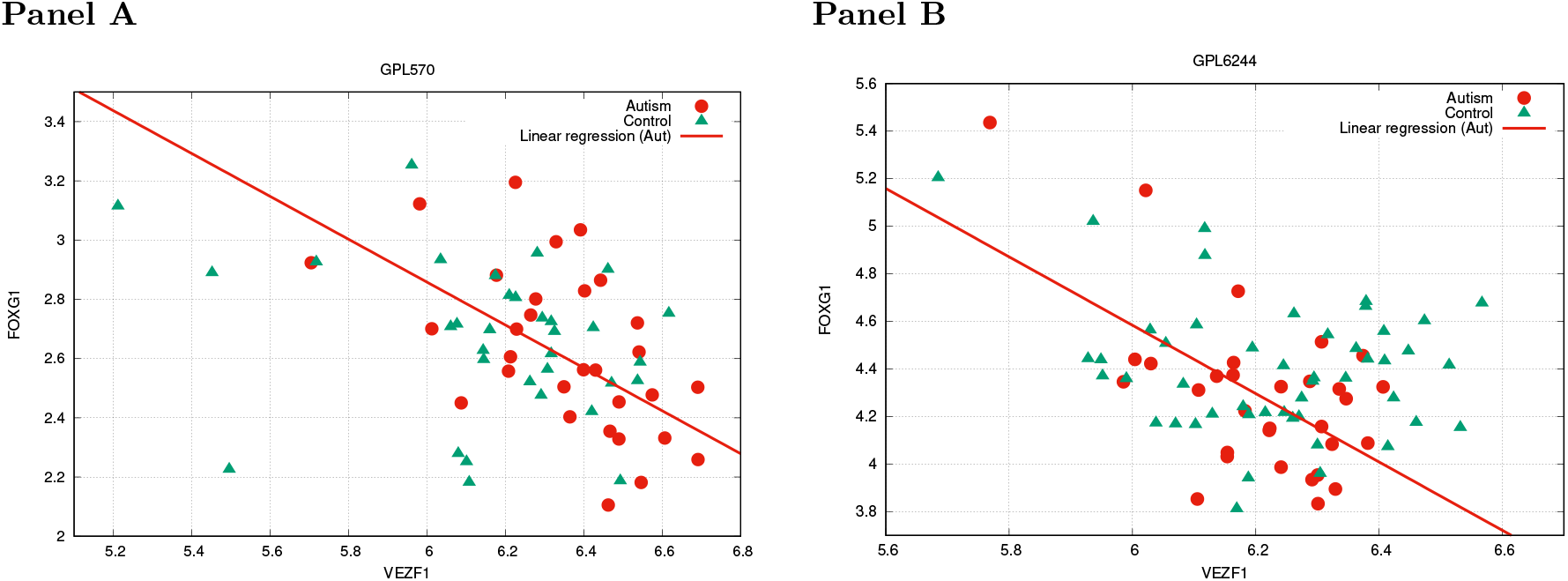
VEZF1 - FOXG1 correlation. VEZF1 expression values (x-axis) are plotted against FOXG1 expression values y-axis). Panel A: red circles are related to GPL570 ASD and green triangles are related to GPL570 controls. Panel B: red circles are related to GPL6244 ASD and green triangles are related to GPL6244 controls. The red lines represent the linear regression associated with ASD samples.

As reported in Table 2 five and ten couples were identified respectively for syndromic and non-syndromic. Interestingly, excluding two couples YY2-MED13 and KLF4-NLGN4X, for all the other couples (13 out of 15) significant positive or negative correlations occur in the ASD while no significant correlation is observed in Controls. It can be hypothesized that the differential and likely altered regulation of those genes, mediated by the corresponding TFs, can be associated with some mechanism triggered by the disease.

## 4 Discussion

In this work, we have studied the differential regulation, mediated by TFs, of key genes involved in ASD. Nine and seven TFs have been identified as potential regulators of the set of syndromic and non-syndromic key high confident genes retrieved by SFARI database. We have also identified significant Transcription Factor - Target Gene couples potentially associated with an altered regulation in ASD patients. Interestingly many identified couples are involved in processes associated with brain morphogenesis and development. Specifically the TF VEZF1 regulates a key brain-specific TF (i.e. FOXG1) involved in brain morphogenesis [18]. This regulation seems to be abrogated in ASD. In the same way, the brain specific TF FOXP2 regulates the TF MEF2C [19], involved in neurogenesis in mouse model, and this association is also abrogated in ASD. Speculatively, the loss of regulation of these brain-specific TFs in ASD, that in turn regulates the transcription of multiple genes involved in brain development, is suggestive of a role of them in the development of ASD.

Less straightforward is the interpretation of the ASD specific regulation of ACTB (the gene that codes for beta actin). ACTB is a housekeeping structural gene with multiple roles, but, interestingly, specific ACTB mutations are linked to deficit in neurodevelopment [20]. Moreover, ZNF148 (whose association with ACTB is significant in ASD) is a zinc finger DNA-binding proteins of the Kruppel family whose mutations have been involved in brain development delay [21, 22]. Please note that the control by VEZF1 on the DYNC1H1 gene that codes for a subunit of Dynein suggests that cytoskeleton dynamics can play a role in ASD. This consideration is supported by the same ZNF148 that controls DPYSL2, that codes for a protein involved in neuron guidance, growth and polarity by promoting microtubule assembly and it has been implicated in multiple neurological disorders [23].

MED13 codes for a subunit of the mediator complex. The mediator complex plays multiple roles in regulating the transcription process, and therefore it is difficult to dissect a specific role for it in ASD [24, 25], even more because it has a negative correlation on non-ASD specimens with YY2 (i.e. the Transcription Factor Yin Yang 2) that can, as the name suggests, acts either as activator or repressor of transcription. Noteworthy, YY2 is also a DNA-binding proteins of the Kruppel family.

A potential role of RNA transcription and regulation is suggested by the presence of CELF4 that code for a brain specific RNA-binding protein involved in the regulation of the splicing [26], under the control of the same VEZF1 discussed above and negatively correlated in ASD. The same CELF4 is also under the control of ZNF263 (and negatively correlated in ASD) that in turn regulates another neuronal specific RNA binding protein (i.e. ELAVL3) and GABR2 that code for a subunit of the GABA receptor, involved at multiple level in ASD, a key component at the synapses the of GABAergic neurons.

At least, the role of TF KLF4, another TF of the Kruppel family, seems central. It is expressed in neural stem cells and controls stemness and regeneration at multiple levels [27]. It positively regulates in ASD ARF3, involved in Golgi trafficking, GIGYF2, involved in the regulation of tyrosine-kinase receptor activities, and PPP5C, that codes for a serine-threonine phosphatase. It also positively regulates NLGN4X (in non-ASD specimens), that codes for a neuroligin, a neuronal cell surface protein involved in the formation and remodeling of central nervous system synapses.

Overall data suggest an involvement of a core network of TFs that regulates brain development and neuronal fate and activities by modulating cytoskeleton dynamics, RNA metabolism and key membrane-bound receptors and related activities.

It is worth noting that obtained computational results, in particular the identified Transcription Factor - Target Gene couples, would deserve further investigations at an experimental level. In this regard, this biased regulation could be the target of some experimental design in order to 1) test this hypothesis and 2) try to target this altered regulation pattern in ASD samples.

It is possible to validate the role of identified transcription factor–target gene pairs in autism by comparing TF binding and target gene expression in cellular models of ASD and controls, using techniques such as coupled ChIP-seq and RNA-seq. Furthermore, exploring whether targeted repression of aberrant TF activity—via CRISPR interference or small molecule inhibitors—can normalize gene expression profiles may elucidate a potential therapeutic strategy.

In conclusion, we would like to emphasize that the present work proposes an effective and reliable computational approach that could be applied to any disease with known key genes and available gene expression data.

## Supporting information

STAB1

## Acknowledgment

This work was carried out as part of the Short Term Mobility (STM) 2023 programme financed by the National Research Council (CNR) of Italy. Daniele Santoni is a member of the Gruppo Nazionale Calcolo Scientifico - Istituto Nazionale di Alta Matematica (GNCS-INdAM). This work is also funded by “BIOSYS3—Optimization, Models and Algorithms for Bioinformatics and System Biology” project (DIT.AD021.128) of the Institute for System Analysis and Computer Science “Antonio Ruberti”—National Research Council of Italy.

## Footnotes

1 https://gene.sfari.org/

2 https://gene.sfari.org/; database version 2023

